# Non-invasive imaging reveals conditions that impact distribution and persistence of cells after *in vivo* administration

**DOI:** 10.1101/202101

**Authors:** Lauren Scarfe, Arthur Taylor, Jack Sharkey, Rachel Harwood, Michael Barrow, Joan Comenge, Lydia Beeken, Cai Astley, Ilaria Santeramo, Claire Hutchinson, Lorenzo Ressel, Jon Smythe, Eric Austin, Raphael Levy, Matthew J. Rosseinsky, Dave J. Adams, Harish Poptani, B. Kevin Park, Patricia Murray, Bettina Wilm

## Abstract

**Background:** Cell-based regenerative medicine therapies are now frequently tested in clinical trials. In many conditions, cell therapies are administered systemically, but there is little understanding of their fate, and adverse events are often under-reported. Currently, it is only possible to assess safety and fate of cell therapies in preclinical studies, specifically by monitoring animals longitudinally using multimodal imaging approaches. Here, using a suite of *in vivo* imaging modalities to explore the fate of a range of human and murine cells, we investigate how route of administration, cell type and host immune status affect the fate of administered cells.

**Methods:** We applied a unique imaging toolkit combining bioluminescence, optoacoustic and magnetic resonance imaging modalities to assess the safety of different human and murine cell types by following their biodistribution and persistence in mice following administration into the venous or arterial system. **Results:** Longitudinal imaging analyses (i) suggested that the intra-arterial route may be more hazardous than intravenous administration for certain cell types; (ii) revealed that the potential of a mouse mesenchymal stem/stromal cell (MSC) line to form tumours, depended on administration route and mouse strain; and (iii) indicated that clinically tested human umbilical cord (hUC)-derived MSCs can transiently and unexpectedly proliferate when administered intravenously to mice.

**Conclusions:** In order to perform an adequate safety assessment of potential cell-based therapies, a thorough understanding of cell biodistribution and fate post administration is required. The non-invasive imaging toolbox used here can expose not only the general organ distribution of these therapies, but also a detailed view of their presence within different organs and, importantly, tumourigenic potential. Our observation that the hUC-MSCs but not the human bone marrow (hBM)-derived MSCs persisted for a period in some animals, suggests that therapies with these cells should proceed with caution.

## Background

In recent years, biomedical and translational research has focussed on exploring the potential of regenerative medicine therapies (RMTs) to treat a vast number of diseases[1]. A primary safety concern of RMTs, especially if based on stem cells, is their potential to form tumours, due to their proliferative and multi-potential differentiation characteristics[2]. Mesenchymal stem/stromal cells (MSCs) isolated from bone marrow, adipose tissue or umbilical cord are being tested in clinical trials for a range of conditions, but in many cases, preclinical safety data are not available, and the authors fail to report whether the cells cause any adverse effects. Bone marrow-derived MSCs have been used for many years and appear safe[3], but a review of adipose-derived MSCs concluded that while adverse events are rare, they nevertheless do occur, and are likely to be related to underlying health conditions of the patients or administration route[4]. Human umbilical cord-derived (hUC)-MSCs have only recently been introduced in clinical trials, with more than 50% of these initiated within the last 3 years (a summary of registered trials in presented in Additional File 1). hUC-MSCs are less immunogenic than other types of MSCs, which contributes to their attraction as clinical RMTs. However, because of their low immunogenicity in combination with higher proliferative behaviour, these cells may also pose a greater potential risk[5], yet until now, their safety profile has not been robustly assessed. The importance of preclinical safety testing is highlighted by a recent report where a tumour developed in a patient’s spinal cord following intrathecal administration of stem cells[6].

Assessing the safety of cell therapies by tracking their distribution and fate over time after administration can be achieved in preclinical models. Many animal studies use lipophilic membrane dyes, such as PKH26 or CM-Dil, to label the cells, which requires culling of animals at various time points and histological analysis[7–11]. The key flaws of this approach are (1) the detection of false positive cells because lipophilic dyes have the potential to transfer to host cells[12]; (2) very large animal numbers, infringing on the principles of the 3Rs (Replacement, Reduction, Refinement); and (3) the failure to longitudinally monitor the cell fate in each individual animal over time. By contrast, non-invasive imaging technologies have opened up exciting new possibilities for preclinical assessment of the safety of cell therapies by allowing longitudinal *in vivo* cell tracking to monitor cell biodistribution and persistence. Preclinical imaging technologies for cell tracking, some of which have clinical relevance, include magnetic resonance imaging (MRI) to detect cells labelled with superparamagnetic iron oxide nanoparticles (SPIONs), multispectral optoacoustic tomography (MSOT) to detect cells labelled with gold nanorods (GNRs) or near-infrared red fluorescent protein[13–17], and bioluminescence imaging (BLI) for the detection of cells expressing the genetic reporter, firefly luciferase[18–20]. Genetic reporters are particularly advantageous because signals are only generated from living cells, thus allowing the monitoring of cell proliferation and tumour growth, and avoiding problems based on nanoparticle dissociation from cells, which can lead to false positive signals. However, the spatial resolution of BLI is poor, making it difficult to precisely locate the cells[18]. By contrast, both preclinical MSOT and MRI have much higher spatial resolution (150 μm and 50 μm, respectively), providing details of the inter- and intra-organ distribution of administered cells. Moreover, as MRI is routinely used in the clinic, it provides a bridge for preclinical and clinical studies.

An advanced approach to longitudinal *in vivo* cell tracking is the use of multi-modal imaging strategies that combine cell labels and reporters, including dual-labelling with both the luciferase reporter gene for BLI, and either gold nanorods (GNRs) for photoacoustic imaging[21–24], or superparamagnetic iron oxide nanoparticles (SPIONs) for MRI[25–28]. Such multi-modal imaging approaches benefit from the sensitivity of the luciferase-based signal conferred by living cells, in combination with the high resolution of MRI and photoacoustic imaging systems to detect the nanoparticles inside organs, allowing a comprehensive longitudinal analysis of cell fate and safety risks.

The most common way to administer cells systemically in small animals is via the intravenous (IV) route through the tail vein [29], delivering cells directly to the lungs where they are sequestered as a consequence of the pulmonary first-pass effect[30–35]. Previous reports have suggested that IV administered cells labelled with lipophilic dyes bypass the lungs, but this is likely due to false positive staining. For example, in renal regenerative studies, PKH26 dye-labelled IV administered cells have been reported to engraft in injured kidneys and replace damaged renal cells [9–11, 36], but a more recent study using this lipophilic dye in combination with GFP expression shows that while the dye can sometimes be detected in the kidneys, the cells remain trapped in the lungs[32]. These recent findings are corroborated by *in vivo* cell tracking studies which show that after IV injection, transplanted cells predominantly accumulate in the lungs [19, 33, 34], fail to integrate or differentiate into tissue-specific cell types and disappear within 7 days [19, 20, 37].

Although the IV route is also frequently used in clinical trials, administration via the arterial circulation is not uncommon. For instance, clinical trials testing the potential of cell therapies to treat myocardial infarction administer cells into the coronary arteries or left cardiac ventricle[24, 38], while in patients with peripheral artery disease or stroke, intra-arterial injection via the femoral or carotid artery, respectively, is frequently employed[39]. Intra-arterial administration will also lead to systemic distribution to other organs, including the brain, and cells passing through the blood-brain barrier could pose an important safety concern. However, a detailed analysis of cell fate after intra-arterial cell administration has so far not been reported [4].

Here, we have implemented a multi-modal imaging approach comprising BLI, MSOT and MRI, to assess biodistribution and fate of different cell types following venous and arterial administration in healthy mice. Some of these cell types are currently being used in clinical trials, including hUC-MSCs (Additional File 1), hBM-MSCs[40], kidney-derived cells[41] and macrophages[42]. We show that our multi-modal imaging approach allows us to determine the immediate distribution of the cells with respect to the route of administration, and to assess the long-term fate of mouse and human MSCs, and their propensity to form tumours. Our findings demonstrate that the multi-modal imaging platform allowing longitudinal cell tracking is an important tool to identify safety concerns of cells used in clinical trials.

## Methods

### Animals

Mice (Charles River, UK) were housed in individually ventilated cages under a 12 hour light/dark cycle, with *ad libitum* access to standard food and water. All animal experiments were performed under a licence granted under the UK Animals (Scientific Procedures) Act 1986 and were approved by the University of Liverpool ethics committee. Experiments are reported in line with the ARRIVE guidelines. Tumour formation was closely monitored and the tumour burden was not allowed to exceed the recommended size[43].

### Cell Preparation

Mouse kidney-derived stem cells (mKSCs)[44], the D1 mouse MSC (mMSC) line (D1 ORL UVA [D1](ATCC^®^ CRL-12424™)), primary human umbilical cord-derived MSCs (hUC-MSCs; collected from consenting donors and produced identically to those already being used in clinical trials by NHS Blood and Transplant (NHSBT)), primary human bone marrow-derived MSCs (hBM-MSCs; Lonza PT-2501), human kidney cells (hKCs; kidneys deemed unsuitable for transplantation via UK NHSBT[32]) and RAW264.7 macrophages (European Collection of Authenticated Cell Cultures 91062702) were cultured at 37°C under a humidified atmosphere with 5% CO_2_ (culture media are described in Additional File 2). Primary human cells were used up to passage 8, whereas mouse lines were cultured up to passage 25.

For detection by BLI, cells were transduced with a lentiviral vector encoding either firefly luciferase (Luc) or a bicistronic construct of Luc and ZsGreen, all under control of the constitutive promoter EF1a. The vector plasmids were a gift from Bryan Welm (Addgene plasmids # 21375 and 39196) and the production of viral particles and cell transduction was carried out as previously described[45,46]. The mKSCs were infected with a multiplicity of infection (MOI) of 10, whereas all other cells were infected with an MOI of 5. At least 90% of the cell populations expressed the vector after transduction, except for macrophages, which did not tolerate polybrene and thus displayed a reduced infection efficiency. Cell sorting based on ZsGreen fluorescence obtained a macrophage population that was 100% positive for the luciferase construct. Cells for karyotyping were treated with colcemid (0.1 μg/mL) followed by a hypotonic treatment and fixation in Carnoy’s fixative. Chromosome analyses were carried out by cytogenetics specialists (CellGS, Cambridge, UK).

Average cell diameter was estimated by measuring the volume of a cell pellet in a packed cell volume (PCV) tube according to the manufacturer’s instructions (Techno Plastic Products, Switzerland). The cell diameter was calculated using the formula:

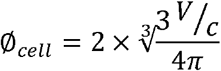

where V corresponds to the pellet volume, and c to the number of cells in the pellet.

For MR tracking, cells were labelled with diethylaminoethyl-dextran coated SPIONs synthesised in house as previously described[25,26]. SPIONs were added to the culture medium at a concentration of 25 μg[Fe]/mL 24h prior to the experiment, after which cells were washed to remove excess particles and harvested for administration as described below. This resulted in an iron content of ~ 6 pg[Fe]/cell.

GNRs were synthesised using a protocol first reported by El-Sayed’s group[47] and coated with silica as described by Comenge *et al.*[23]. Macrophages were labelled for 24h with GNRs at a final concentration of 10 pM before harvesting for cell injection. Neither of the labelling approaches (SPIONs/GNRs) caused any reduction in cell viability.

### Cell administration

Cells were trypsinised, pelleted, resuspended in ice-cold phosphate buffered saline (PBS) and kept on ice until injection. 100μl cell suspension was administered to mice via intravenous (IV) or ultrasound-guided intracardiac (IC) injection. A description and comments on this method of administration is provided in Additional File 3.

### Bioluminescence imaging

Short-term study: ZsGreen^+^/Luc^+^ mMSCs, mKSCs, hKCs, macrophages or Luc^+^ hUC-MSCs were administered IV or IC to BALB/c mice. Long-term study: ZsGreen^+^/Luc^+^ mMSCs or Luc^+^ hUC-MSCs were administered by IV or IC to BALB/c (severe combined immunodeficient) SCID mice (see Table 1 for route, cell dose and number of animals in each experiment). The *in vivo* biodistribution of cells was monitored by BLI immediately after cell administration, and at multiple time points up to 30 days. Mice were administered 150 mg/kg body weight luciferin (Promega, UK) subcutaneously, and imaged 15 min later in a bioluminescence imager (IVIS Spectrum, Perkin Elmer, UK). Imaging data were normalised to the acquisition conditions and expressed as radiance (photons/second/cm^2^/steradian (p/s/cm^2^/sr)), and the colour scale was adjusted according to the strength of signal detected. Because IV injections into the tail can lead to a small fraction of cells remaining in or around the injection site, causing strong signal intensities, the tails of animals that received cells via this route were covered prior to data acquisition. At the respective study end points, mice were culled and organs with any visibly identifiable tumours imaged *ex vivo* by BLI. Kidneys were cut coronally for *ex vivo* imaging, and all other organs were imaged whole. Bioluminescence signals of whole live mice or individual organs ex vivo were quantified by drawing regions of interest (ROIs) from which the total flux (photons/second) was obtained. The relative signal intensity from each organ was calculated as a percentage of the signal intensity from all organs. For *ex vivo* kidney imaging, the ROI was drawn around all four kidney halves and a single value for total bioluminescence signal was recorded.

**Table 1.** Experimental details of studies, mouse strains, cell types, route of administration, dose and number of animals studied

| Study | Mouse strain | Cell Type | Route | Dose | Number of animals |
| --- | --- | --- | --- | --- | --- |
| BLI short-term biodistribution (and MSOT for RAW macrophages) | BALB/c | mMSC | IV and IC | $1.0 \times 10^6$ | Minimum n = 3 for each cell type and administration route |
| | BALB/c | mKSC | IV and IC | $1.0 \times 10^6$ | |
| | BALB/c | hUC-MSC | IV and IC | $1.0 \times 10^6$ | |
| | BALB/c | hBM-MSC | IV and IC | $5.0 \times 10^6$ | |
| | BALB/c | hKC | IV and IC | $0.3 \times 10^5$ | |
| | BALB/c | macrophages | IV and IC | $1.0 \times 10^7$ | |
| BLI long-term biodistribution | BALB/c SCID | mMSC | IV | $1.0 \times 10^6$ | n = 3 |
| | BALB/c SCID | mMSC | IC | $1.0 \times 10^6$ | n = 5 |
| | BALB/c | mMSC | IC | $1.0 \times 10^6$ | n = 4 |
| | FVB | mMSC | IC | $1.0 \times 10^6$ | n = 4 |
| | MF1 | mMSC | IC | $1.0 \times 10^6$ | n = 4 |
| | BALB/c SCID | hUC-MSC | IV | $1.0 \times 10^6$ | n = 13 |
| | BALB/c SCID | hUC-MSC | IC | $1.0 \times 10^6$ | n = 3 |
| | BALB/c SCID | hUC-MSC | IV | $5.0 \times 10^5$ | n = 6 |
| | BALB/c SCID | hUC-MSC | IC | $5.0 \times 10^5$ | n = 14 |
| | BALB/c SCID | hBM-MSC | IV | $5.0 \times 10^5$ | n = 6 |
| | BALB/c SCID | hBM-MSC | IC | $5.0 \times 10^5$ | n = 6 |
| MRI cell | BALB/c | mMSC | IV | $1.0 \times 10^6$ | n = 2 |
| tracking | BALB/c | mMSC | IC | $1.0 \times 10^6$ | $n = 7$ |

### Multispectral Optoacoustic Tomography (MSOT)

MSOT was carried out using the inVision 256-TF MSOT imaging system (iThera Medical, Munich). Images were recorded at the following wavelengths: every 10 nm from 660 nm and 760nm, and every 20 nm from 780 nm and 900 nm, at a rate of 10 frames per second and averaging 10 consecutive frames. All mice were allowed to equilibrate in the imaging system for 15 minutes prior to recording data. For monitoring of the biodistribution of macrophages after IV administration, a 15 mm section of the abdomen to include the liver, kidneys and spleen of the mice was imaged repeatedly for a total of 4.5 hours; 30 minutes into the imaging the mice received the macrophages via a tail vein catheter. For the IC imaging a 15 mm section of the abdomen was imaged once, followed by an ultrasound (Prospect 2.0, S-Sharp, Taipei city) guided injection of 10^7^ macrophages into the left ventricle of the heart. Mice were then returned to the photoacoustic imaging system for imaging as previously described. Data was reconstructed and multispectral processing was performed to resolve signals in the liver, kidney and spleen for GNRs. Regions of interest were drawn around the liver, right kidney and spleen (an example is shown in Additional File 4) to generate mean pixel intensity data.

### MR imaging

ZsGreen^+^/Luc^+^/SPION^+^ mMSCs (10^6^) were administered to BALB/c mice IV (n = 2) or IC (n = 2 for short-term analysis; *n* = 5 for longitudinal tracking). The biodistribution of cells in the brain and kidney was imaged with a Bruker Avance III spectrometer interfaced to a 9.4T magnet system (Bruker Biospec 90/20 USR) using a Fast Low-Angle Shot (FLASH) T_2_* weighted sequence at baseline and up to 2 days post administration. T_2_* relaxation times were obtained from a T_2_* map generated with a multi-gradient echo sequence by drawing ROIs around the cortex of the kidney (an example is shown in Additional File 5) or a region of the liver. At least one animal was culled at each time point for histological analyses, and brains and kidneys were fixed with 4% formaldehyde and imaged at a higher resolution *post mortem* (all MRI acquisition parameters are described in the Additional File 6). Tumours were imaged with a T_2_ weighted fast spin echo sequence.

### Histopathological analysis

Perfusion fixed frozen brain and kidney sections were stained for the endothelial cell marker isolectin B4 (IB4, L2140, Sigma Aldrich, UK) as described previously[48]. The presence of ZsGreen^+^ mMSCs within brain and glomerular capillaries was imaged by confocal microscopy (LSM 800 Airyscan, Zeiss). Frozen kidney sections (7μm) were stained for the presence of iron (Iron Stain Kit, Sigma, UK) according to manufacturer’s instructions to detect SPIONs, and consecutive sections were counterstained with DAPI. Prussian blue stained cells and ZsGreen-positive mMSCs were imaged by bright field and epifluorescence microscopy.

Tumours were fixed in 4% paraformaldehyde at 4°C for 24h, washed in PBS and processed through an ethanol and xylene series before embedding in paraffin. Five μm tissue sections were stained for haematoxylin and eosin (H&E) by standard methods and morphologically assessed.

### Fluorescence activated cell sorting (FACS)

Bone marrow was extracted as previously described[49]. In short, femurs and tibias were collected in PBS containing penicillin/streptomycin, the bone marrow was flushed out with PBS, centrifuged (400 g, 5 mins) and then resuspended in fresh PBS before analysis by flow cytometry for ZsGreen expression.

### Statistical Analyses

Statistical analyses were performed using Minitab 17 statistical software. A one-way analysis of variance (ANOVA) was used to compare multiple groups. When an ANOVA resulted in a statistically significant result (p < 0.05), a Tukey pairwise comparison was performed in order to determine which groups were significantly different. The Tukey pairwise comparison assigned each group at least one letter, and groups that did not share a letter were significantly different from one another.

## Results

### Whole body biodistribution of different cell types following intravenous (IV) and intracardiac (IC) administration

Bioluminescence imaging showed that IV delivery of ZsGreen^+^/Luc^+^ mouse MSCs (mMSCs), mouse kidney-derived stem cells (mKSCs) and human kidney cells (hKCs) resulted in signals exclusively in the lungs, while signals from IV-administered macrophages were also located more posteriorly (Fig. 1a). This was expected because macrophages are known to traverse the lungs and populate other organs, such as the liver and spleen. In contrast, intra-arterial delivery via the left heart ventricle (from now on referred to as intra-cardiac (IC)) resulted in a whole-body distribution of all cell types (Fig. 1a).

**Figure 1.**
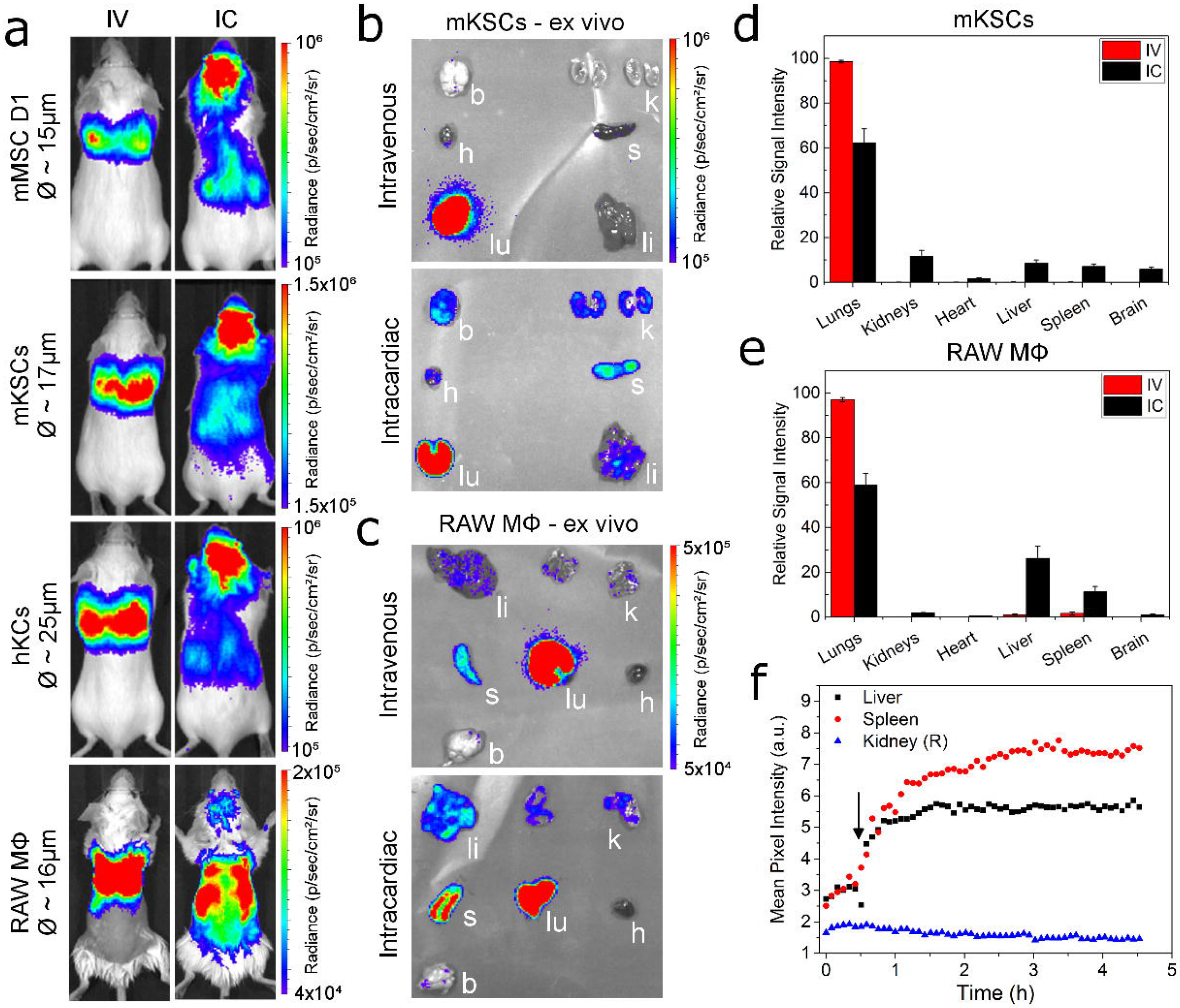
Biodistribution of different cells following intravenous or intracardiac administration. **(a)** BLI immediately after administration, showing that cells were always confined within the lungs after intravenous (IV) administration, but distributed throughout the body after intracardiac (IC) administration; an exception were the macrophages which showed also a more posterior signal after IV administration. The diameter of each cell as estimated by the PCV is shown next to the images. *Ex vivo* bioluminescence imaging of organs within 5h of administration of **(b)** mKSCs or **(c)** macrophages confirmed the *in vivo* cell biodistribution. Organs are indicated as kidneys (k), spleen (s), liver (li), lungs (lu), heart (h) or brain (b) and the colour scale applies to both administration routes. Quantification of the bioluminescence signal intensity of organs *ex vivo* post **(d)** mKSC or **(e)** macrophage administration. Values represent the mean signal intensity measured in each organ and normalised to the total flux from all organs (n = 3 each group). Error bars represent standard error. **(f)** Mean pixel intensity of GNR-labelled macrophages measured via multispectral optoacoustic tomography for a period of 5 hours post IV administration, displaying the kinetics of their accumulation in the spleen and liver. Arrow indicates the time point at which the cells were administered.

Organ-specific *ex vivo* imaging within 1h of IV administration of mKSCs confirmed that the signal was limited to the lungs (Fig. 1b, d). In contrast, after IC administration, bioluminescent signals were detected in the brain, heart, lungs, kidney, spleen, and liver (Fig. 1b, d). IV-administered macrophages were found predominantly within the lungs by *ex vivo* imaging (Fig. 1c), but weaker signals were also detected in the spleen and liver, kidneys and brain, confirming the *in vivo* signal distribution. *Ex vivo* analysis of macrophages after IC injection showed signals in most organs that were imaged (Fig. 1c, e).

To monitor the temporal dynamics of macrophage migration, cells were labelled with GNRs, injected IV, and monitored continuously for 4.5h using MSOT. Signal intensity began to increase immediately in both the liver and spleen until around 90 min when it started to plateau (Fig. 1f), but remained close to basal levels in the kidney, consistent with BLI *ex vivo* analysis (Fig. 1e, f). However, when GNR-labelled macrophages were administered IC, increases in signal intensity in the kidney were comparable to those in the liver and spleen 4h post-administration (quantification is shown in Additional File 4c).

### Cell distribution within organs using high-resolution magnetic resonance imaging

Since the spatial resolution of BLI is poor, we used MRI to evaluate the intra-organ biodistribution of ZsGreen^+^/Luc^+^/SPION^+^ mMSCs after IV or IC administration, focussing particularly on the brain and kidneys. Following IC injection, T_2_* weighted imaging revealed hypointense areas distributed homogenously throughout the brain (Fig. 2a), and localised in the cortex of the kidneys (Fig. 2b). However, hypointense contrast was not detected in the brain or kidneys of IV-injected mice, confirming that IV administration does not deliver mMSCs to either of these organs (Fig. 2a, b). Post mortem MR imaging of extracted organs performed at higher resolution confirmed the hypointense contrast throughout the brain and in the renal cortex of IC-injected mice (Fig. 2a, b).

**Figure 2.**
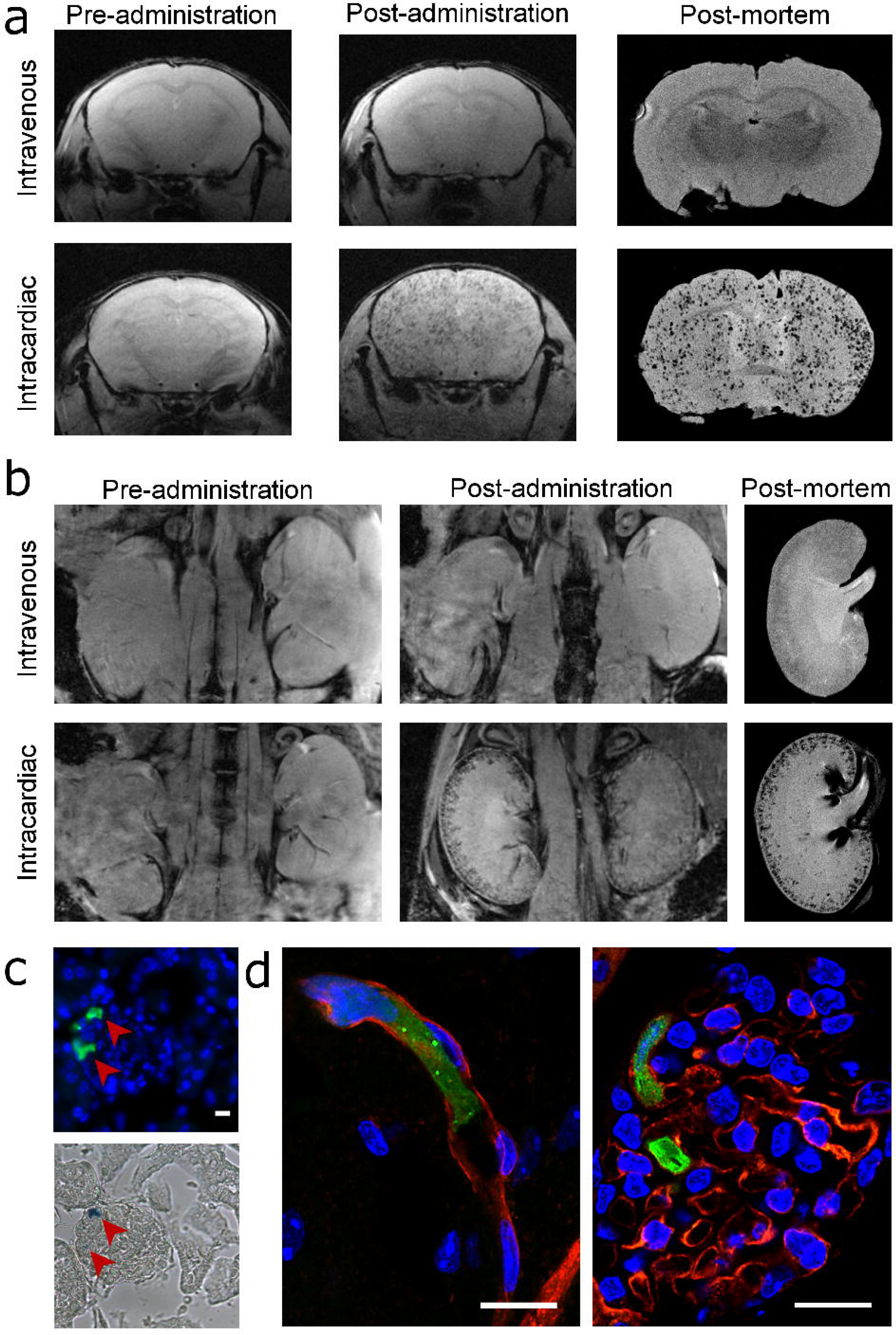
MRI and immunofluorescence images of mMSCs in the brains and kidneys. *In vivo* and post-mortem T_2_* -weighted images of the **(a)** brains and **(b)** kidneys of mice pre- and post-administration of SPION-labelled mMSCs via the IV or IC route. **(c)** Epifluorescence of Zsgreen (green) and nuclei (blue) of a single kidney glomerulus (top) and the corresponding Prussian Blue image (bottom) demonstrating that cells and SPIONs co-localised to the same spatial location. Scale bars correspond to 10 μm. **(d)** Overlay of confocal microscopy images of Isolectin-IB 4 staining (red), ZsGreen (green) and nuclei (blue). Tissue sections were obtained from the brain (left) or kidney (right) of mice that received cells IC.

Histological analysis of ZsGreen expression by fluorescence microscopy in combination with Prussian Blue staining of SPIONs showed that labelled cells were located in the renal glomeruli (Fig. 2c). ZsGreen and Prussian Blue signals corresponded to the same spatial location, indicating that hypointense contrast *in vivo* was unlikely to result from false-positive detection of SPIONs (e.g. released from dead cells). To determine whether IC-administered cells had undergone extravasation, we performed confocal imaging of IB4-stained blood vessels. This demonstrated that ZsGreen+ mMSCs were physically trapped in the lumen of microcapillaries (Fig. 2d), suggesting that the cells did not cross the blood brain barrier or the glomerular filtration barrier.

### Short-term fate of IC-injected cells

To determine how long the cells persisted in major organs we injected 10^6^ ZsGreen^+^/Luc^+^/SPION^+^ mMSCs into the left cardiac ventricle of BALB/c mice and tracked their fate *in vivo* by MRI and BLI, and post mortem by MRI and fluorescence microscopy (Fig. 3a). On the day of injection, whole-body distribution of IC-administered mMSCs by bioluminescence signals was observed, while in the kidneys, MRI revealed hypointense contrast specifically in the cortex. By 24h, the bioluminescence signal intensity decreased, suggesting cell death. Correspondingly, fewer hypointense areas were observed in the renal cortex by MRI, supporting the disappearance of SPION-labelled cells. By 48h, bioluminescence was no longer detectable in the abdominal region, nor was any significant hypointense SPION contrast observed in the kidneys with MRI. This was confirmed by high-resolution MRI of organs *ex vivo,* showing a decrease in contrast in the renal cortex over time, and a decrease in the frequency of ZsGreen^+^ mMSCs in kidney glomeruli by fluorescence microscopy (Fig. 3a). Changes in the T_2_* relaxation time in the renal cortex indicated the relative number of SPION-labelled cells present at each time point. T_2_* was significantly lower on the day of cell administration (Fig. 3b) than at baseline but then increased towards baseline levels at 24h and 48h. Because the liver is the major organ for clearance of blood-transported particulates, we quantified the hepatic T_2_* relaxation time, which revealed a subtle but significant decrease from baseline through to 48h (Fig. 3c). These results suggest that following cell death, SPIONs accumulate predominantly in the liver and are not retained by the kidneys.

**Figure 3.**
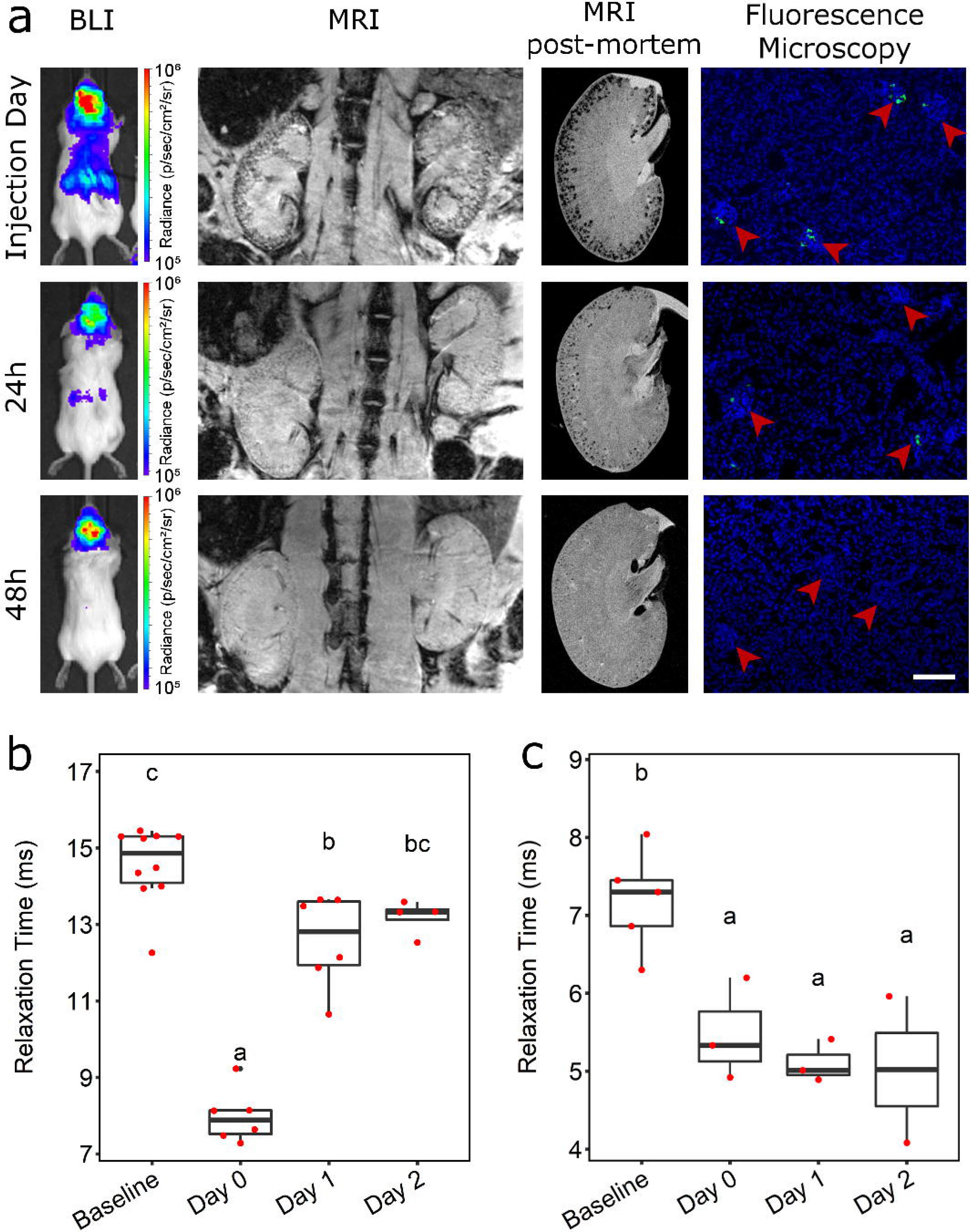
Short-term fate of mMSCs imaged *in vivo* and post-mortem. **(a)** BLI, MRI (*in vivo,* post-mortem) and fluorescence microscopy images of the kidneys immediately, 24h or 48h after IC administration of SPION-labelled mMSCs. Fluorescence images were obtained from tissue sections where green fluorescence corresponds to ZsGreen expression and blue fluorescence to DAPI staining. Arrowheads indicate individual glomeruli. Scale bar corresponds to 100 μm. T_2_* relaxation time of **(b)** kidney cortices or **(c)** liver before (baseline) and up to 2 days after cell administration. The T_2_* relaxation time in the cortex of the kidney was significantly lower on the day of cell administration (day 0, mean = 7.98 ms +/− SE = 0.29) than at baseline (14.56 +/− 0.32 ms; One-way ANOVA, p < 0.001). The T_2_* relaxation time then increased towards baseline levels at day 1 (12.57 +/− 0.50 ms) and day 2 (13.19 +/− 0.23 ms), and by day 2 the difference compared with baseline levels was no longer statistically significant. In the livers, T_2_* relaxation time revealed a subtle but significant decrease in relaxation time from baseline to day 2 (baseline, 7.19 +/− 0.29 ms; day 0, 5.48 +/− 0.38 ms; day 1, 5.10 +/− 0.16 ms; day 2, 5.02 +/− 0.94 ms; One-way ANOVA, p = 0.006). Time points that do not share the same letters are significantly different from one another, p < 0.05 (Tukey’s post hoc test).

### Effect of administration route on the long-term biodistribution and fate of mMSCs

To assess the effect of administration route on the long-term fate of cells, ZsGreen^+^/Luc^+^ mMSCs were administered to BALB/c SCID mice by IC or IV routes, and biodistribution monitored by BLI at multiple time points over 28 days. While both IC and IV injection resulted in the typical immediate biodistribution patterns by 24h (Fig. 1a), by 96h following IV and IC administration, the bioluminescence signal was undetectable, indicating loss of cells via cell death (Fig. 4a). Continued imaging over time showed that bioluminescence signals began to increase again in animals after IC injection from around day 14, but not in animals after IV injection. The increase in signal was particularly prominent in the hindquarters of all five IC-injected mice at day 14, and increased further until day 30 (Fig. 4a, Additional File 7a). Detailed analysis of animals after IV administration of mMSCs revealed that bioluminescence signals in the lungs of one mouse increased over time (Additional File 7b). Overall, whole-body bioluminescence intensity initially decreased following both IC and IV administration, and subsequently increased rapidly in the IC-injected mice (Fig. 4b-d).

**Figure 4.**
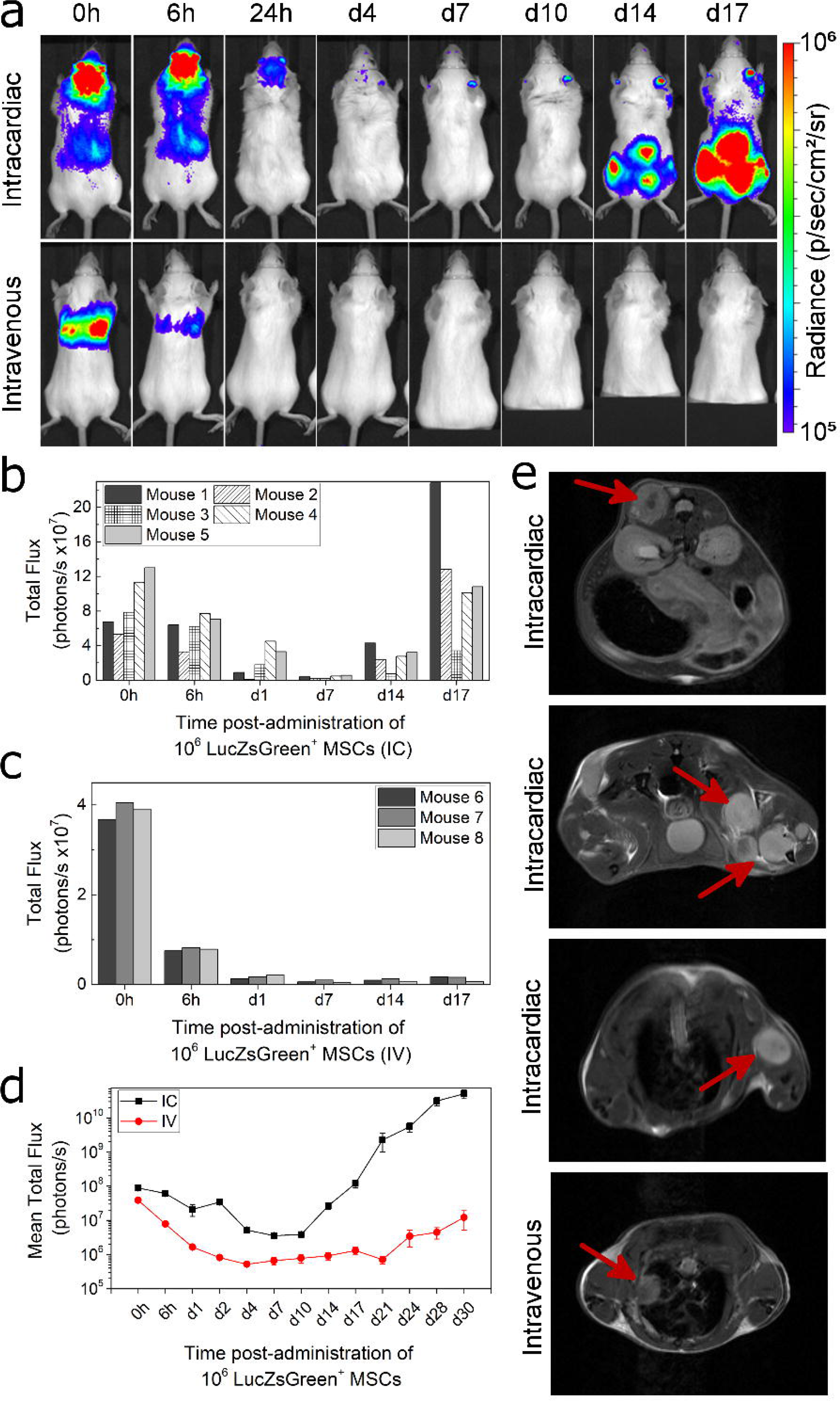
Impact of administration route on long-term tumour formation. **(a)** Representative BLI of SCID mice administered with mMSC via the IC or IV route. BLI scale corresponds to levels between 1.0×10^5^−1.0×10^6^ p/s/cm^2^/sr. Quantification of the bioluminescence signal from each individual mouse that received mMSCs **(b)** IC (n=5) or **(c)** IV (n=3) up to day 17. Signal corresponds to a region of interest drawn around the whole body of the mouse. **(d)** Mean whole body quantification of the bioluminescence signal up to day 30. Error bars represent SE. **(e)** T_2_-weighted MRI of tumours in animals that received mMSCs via IC or IV as imaged 30 days post administration. Arrows indicate individual tumours, usually in the skeletal muscle apart from the IV route, where a tumour was found close to the lungs.

### Osteosarcoma formation after IC administration of mMSCs

Multiple abnormal growths were present in IC-injected BALB/c SCID mice, predominantly in skeletal muscle surrounding the femurs, but also in muscle near the hips, ribs, and spine (Fig. 5a, f), suggesting tumours had formed. Tumour sites corresponded to foci of intense BLI signals which could also be identified using T_2_ weighted MR imaging (Fig. 4e). Furthermore, T_2_ weighted MR imaging allowed us to detect an abnormal mass in the lungs of one (out of three) IV-injected mouse that displayed an intense bioluminescence signal (Fig. 4e, Additional File 7b). Although cells of the mMSC line have been suggested to home to the bone marrow[50], flow cytometry analysis showed the bone marrow was negative for ZsGreen+ cells (Additional File 8). Histologically, tumours were characterised by atypical solid proliferation of spindle cells associated with multifocal formation of pale amorphous eosinophilic material (osteoid). The tumours were therefore classified as osteosarcomas (Fig. 5h, j, k). Frozen sections of the tumour tissue exhibited specific ZsGreen fluorescence (Fig. 5i), further confirming the neoplasms originated from mMSCs.

**Figure 5.**
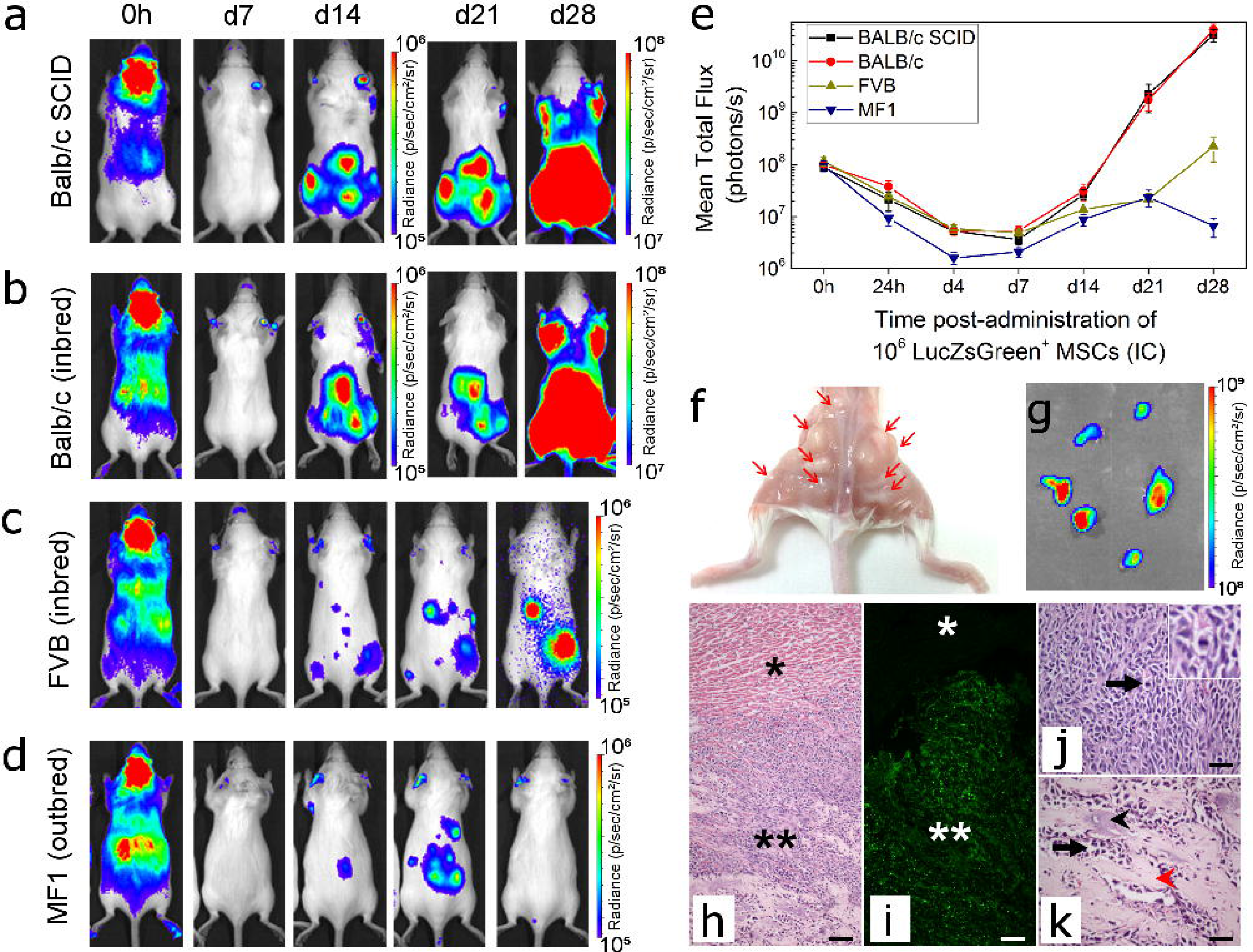
Tumour-formation potential in different mouse strains. Representative BLI of longitudinal tumour monitoring in four strains of mouse following IC administration of mMSCs. **(a)** immunocompromised BALB/c SCID, **(b)** Immunocompetent BALB/c **(c),** FVB or **(d)** MF1 mice. BALB/c mice showed very similar tumour formation potential to BALB/c SCID mice, with respect to timing, size and location of tumour development. After 21 days, the strong signal originating from the tumours required a colour scale two orders of magnitude greater than that at 0h to accurately display the tumour location. FVB and MF1 mice displayed weaker BLI foci at d28, and not all animals displayed the same tumour distribution. Balb/c SCID data in **(a)** has been partially reproduced from Fig. 4a to facilitate comparison between strains. **(e)** Mean whole body quantification of the bioluminescence signal up to day 30. Error bars represent SE. **(f)** Photograph of the hindquarters of a Balb/c mouse after removal of the skin. Multiple tumour foci are indicated with arrows, demonstrating their presence in the skeletal muscle close to the femurs, hips, and spine. **(g)** BLI of tumours harvested 30 days post-administration of mMSCs confirming that tumours originated from the administered cells and not host tissue. **(h-k)** Histological examination of tumour tissue. **(h)** H&E staining and corresponding (**i**) epifluorescence imaging of the ZsGreen reporter. Differences in cell composition between the tumour (**) and normal tissue (*) are denoted. Scale bars represent 100 μm. **(j, k)** Higher magnification of tumour tissue showing cancer cells arranged in densely cellular monomorphic areas. Scale bars correspond to 50 μm and arrow indicates mitotic figures, one of which is shown in the inset. **(k)** Corresponds to an area where the tumour is moderately cellular with production of unmineralised osteoid (black arrowhead) and partially mineralised matrix (red arrowhead).

Chromosomal analysis of the mMSCs revealed a grossly abnormal karyotype of between 65 and 67 chromosomes, with multiplications and unidentified chromosomes (shown in Additional File 9a).

### Formation of mMSC-derived tumours in different mouse strains

To determine whether tumours developed because the BALB/c SCID mice were immunocompromised, we investigated the long-term fate of the mMSCs following IC administration in three different immunocompetent mouse strains: BALB/c (same genetic background as mMSCs), FVB (unrelated inbred strain), and MF1 (unrelated outbred strain). The biodistribution immediately after injection was similar between the strains, but at day 28, only the BALB/c mice displayed bioluminescence signals as high as those in the BALB/c SCID mice (Fig. 5a-d). Moreover, the timing and location of tumour formation was consistent in all immunocompetent and immunocompromised BALB/c mice. In the FVB and MF1 strains, mMSC foci tended to form in similar locations as with the BALB/c mice, but bioluminescence signals were weaker. Although signal intensity gradually increased in FVB mice from d7 to d28, in MF1 outbred mice, signals increased initially up to d21, but then started to decrease as the mMSC foci began to regress (Fig. 5e).

### Long-term biodistribution of hUC-MSCs in BALB/c SCID mice

Since mMSCs gave rise to tumours in immuno-compromised and -competent mice, predominantly after systemic arterial injection, we aimed to determine whether clinically relevant MSCs could carry a similar health risk. We focussed on well-studied hBM-MSCs as well as hUC-MSCs, as the latter are currently being used in an increasing number of clinical trials (Additional File 1). The chromosomal analysis for both hBM-MSCs and hUC-MSCs revealed a normal karyotype (Additional File 9b,c).

When following the fate of both of these cell types after IV or IC administration in BALB/c SCID mice, we found that in most cases, BLI signals became weaker within a few days of administration, and remained undetectable for the duration of the study (hBM-MSCs: 4 weeks; hUC-MSCs: 8 weeks) (Fig. 6a). Ex vivo analysis of the organs on the day of injection suggested that similar to the other cell types, a whole-body distribution is obtained when cells are injected into the arterial system, and cells are mostly trapped in the lungs when the venous route is used. However, in the case of hUC-MSCs, BLI signal was sometimes observed in the heart (Figure 6b). When imaging the same organs without the lungs, and with an increased detection sensitivity, the signal in the heart became more obvious, while very weak signals could also be observed in other organs (Additional File 10). Interestingly, long-term imaging of mice that had received hUC-MSCs via the IV route revealed that in a small number of animals (~25%) foci had developed in locations beyond the lungs (Fig. 6c, red arrows), although in all cases these regressed within the time course of the experiment.

**Figure 6.**
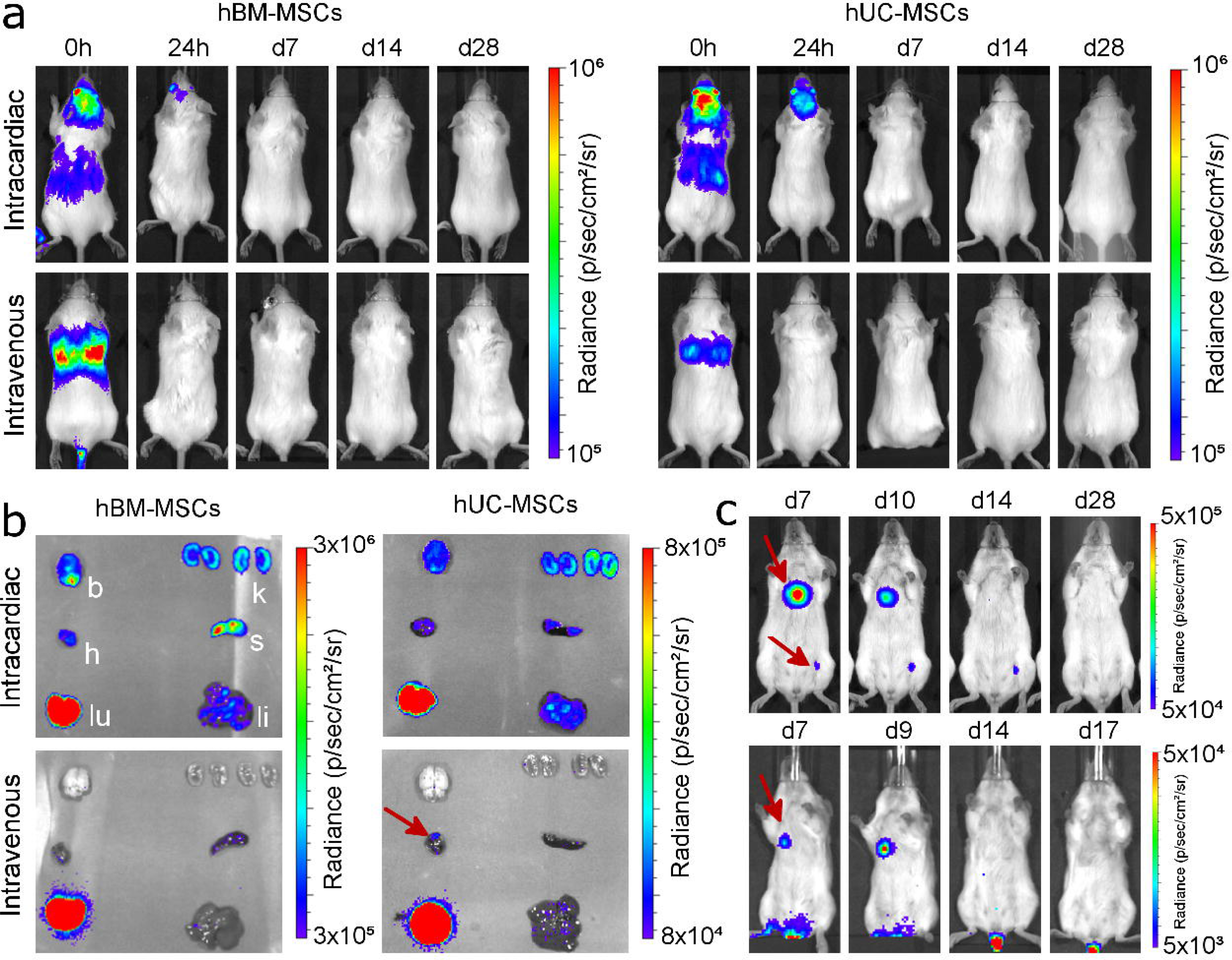
Long-term monitoring of human MSCs in Balb/c mice. **(a)** Representative BLI of mice administered with 5×10^5^ hBM-MSC or hUC-MSC via the IC or IV route. The signal was progressively lost shortly after administration, with no evidence of malignant growth. **(b)** Ex vivo bioluminescence imaging of organs within 5h of administration of the cells. Organs are indicated as kidneys (k), spleen (s), liver (li), lungs (lu), heart (h) or brain (b). In some occasions signal foci were seen in the heart of mice that received hUC-MSC IV (red arrow). **(c)** BLI images from mice that displayed hUC-MSC signal that persisted beyond day 7 (ventral orientation, lower scale). In all cases, the signals had disappeared by day 21 and had not returned by the end of the experiment.

## Discussion

Here, we have employed a novel platform approach of non-invasive preclinical imaging encompassing BLI, MRI and MSOT to assess the biodistribution and persistence of a range of mouse and human cell types following IV and IC administration in healthy mice. These cells included mouse MSCs, kidney stem cells and macrophages, as well as human kidney-derived cells and two types of human MSCs, which are already being tested as cell therapies in clinical trials. As expected, immediate analysis after IV administration revealed that apart from macrophages, all other cell types were mostly sequestered in the lungs, although small numbers of hUC-MSCs could be detected in other organs following *ex vivo* analysis. After IC administration, all cell types showed a widespread distribution. However, irrespective of the administration route, analysis using all three imaging technologies determined that cells disappeared from major organs within 24-48 hours, which based on the loss of BLI signals, was likely due to cell death. The observation that cells are cleared very quickly from the major organs following IC administration indicates that the arterial route poses no significant advantage for cell therapy administration. However, it is possible that small numbers of cells reaching the organs after IC administration may still be able to locally exert beneficial roles through paracrine effects before they are cleared. Therefore, further research is required to explore the local role of IC delivered cells.

Our platform of imaging techniques was also able to provide some mechanistic insight into the fate of cells after administration. Macrophages have been previously shown to home to the liver and spleen after passage through the lungs[51]. However, the dynamics of this homing process had not been described. Using multi-modal BLI and MSOT, we could monitor macrophage accumulation in the liver and spleen for 4.5h continuously at high spatial resolution. We found that labelled macrophages immediately started to accumulate in liver and spleen, particularly in the first ~90 min, which indicated that some of the macrophages instantly passed through the pulmonary circulation.

While BLI has the advantage of highly sensitive body-wide detection of luciferase-expressing cells, its spatial resolution is poor, which prevents organ-focussed imaging. To visualise cells within major organs such as kidney and brain, and monitor their fate over time, we implemented a bimodal approach comprising BLI and MRI, taking advantage of the high spatial resolution of MRI in addition to the high sensitivity of BLI and the fact that luciferase activity is dependent on cell viability[15–18]. Detailed analysis of the biodistribution of mMSCs after IC injection using *in vivo,* and subsequently *ex vivo* MR imaging techniques revealed that SPION-labelled cells were scattered throughout the brain, while in the kidneys, they were restricted to the cortical regions. *Ex vivo* histological staining and fluorescence microscopy demonstrated that cells in the kidneys were found only within the glomeruli, bounded by endothelial cells within the microvasculature, where they appeared to be trapped. Similarly, cells in the brain were only localised within the microvasculature, indicating that they lack the capacity to pass through the blood brain barrier. These results demonstrate that the mMSCs cannot extravasate into the brain and kidneys.

During long-term cell tracking of the BALB/c-derived mMSCs, we observed tumour formation in skeletal muscle following IC administration to a similar degree in immune-competent BALB/c mice as in BALB/c SCIDs. mMSCs also gave rise to tumours in an unrelated inbred strain, albeit at a slower rate, while in an unrelated outbred strain, small foci of mMSCs expanded at early time points and later regressed. Taken together, these data suggest that the adaptive immune system might not be able to recognise tumours derived from syngeneic MSCs, and that the genetic background of the host appears to have an effect on the propensity of MSCs to form tumours. This could be a concern for human trials using autologous MSCs where the ability of the cells to form tumours may not be detected by the recipient’s immune system. Furthermore, the results suggest that the risk of tumour formation might depend on undefined genetic factors that would vary from patient to patient. However, it is important to note that karyotype testing of the mMSC line used here revealed a range of chromosomal abnormalities, which could contribute to their propensity to form tumours.

Our observation that mMSCs distributed to most organs following IC injection, but tumours were predominantly localised in the skeletal muscles and not within the organs they originally appeared in, raises the question of how tumour formation is regulated in different organs and tissues. Our data indicate that the cells had a ‘survival advantage’ in muscular tissue, but not in the brain and the kidneys, from which they failed to extravasate. We hypothesise that following IC administration, a small number of MSCs were able to extravasate from the capillaries in the skeletal muscle where they started to proliferate. The mechanisms that regulate the ability of the mMSCs to extravasate and form tumours in the skeletal muscle but not in other organs are not known, and further analysis is required to determine the molecular and cellular factors controlling this process.

Our results also show that the cells failed to home to and populate the bone marrow, which is surprising given the cells had been originally isolated from the bone marrow[52]. The D1 mMSC line used here has not previously been reported to generate invasive tumours, since subcutaneously injected cells provided no evidence of metastasizing, even if they proliferated at the injection site[23,53]. Our observation that the mMSCs did not form tumours outside the lung following IV administration is therefore consistent with this finding. However, the formation of osteosarcomas in the skeletal muscle after IC administration of mMSCs in this study is in line with the previously described formation of osteosarcomas after adoptive transfer of primary MSCs, particularly for cells expanded *in vitro,* and is a major safety concern in therapies using MSCs[54,55].

Since these observations suggested that arterial administration of MSC-based cell therapies could have important safety implications, we followed the fate of MSC derived from the bone marrow or the umbilical cord of healthy human donors. We confirmed that neither of these cells presented any major chromosomal aberrations, even after transduction with the luciferase reporter. While in most animals the cells became undetectable within a few days after IV administration, in a few mice the hUC-MSCs persisted longer, albeit transiently, in other body regions where their presence was not expected. Of note, a transient persistence of hUC-MSCs was not observed after IC administration, nor for hBM-MSCs using IV or IC administration routes. We suggest that this unusual behaviour is not linked to cell size, because the hUC-MSCs are not smaller than mKSCs or mMSCs, but could possibly be due to their surface proteins, allowing some of the cells to escape the lungs[33,56]. The observation that hUC-MSC foci appeared in a small number of mice, grew in size, but later disappeared, was difficult to explain, especially given that the mice were SCIDs and thus lacked an adaptive immune system. It is possible that the cells eventually elicited a xenogeneic response involving macrophages and natural killer cells[57], after initially suppressing the native immune system, which is one of their central properties[58,59]. Alternatively, the hUC-MSCs may have expanded in the animal but then become senescent and died, irrespective of the host’s ability to mount an immune response. Thus, after an 8-week period of cell tracking, we could not observe any tumour growth by *in vivo* or *ex vivo* BLI in all SCID mice to which hUC-MSCs had been administered by either IV or IC injection. However, the observation that the IV-injected hUC-MSCs persisted for a time period in 25% of the animals indicates that these cells carry greater safety risks and suggests that clinical therapies with these cells should proceed with caution with an appropriate risk management plan. Further preclinical studies are needed to determine the mechanisms by which hUC-MSCs were able to persist as well as eventually disappear in order to better define the potential for tumourigenicity. The imaging platform presented here provides the necessary biotechnology for preclinical evaluation of the potential tumourigenicity of cell products used for cell transplantation, for which there is presently no internationally recognised guideline.

## Conclusions

Cell-based therapies are currently being considered for a range of diseases, some of which are already undergoing clinical trials. A robust biodistribution and safety assessment is essential for understanding cell fate and ensure patient welfare. Here, we demonstrate a safety assessment toolkit that can expose not only the general organ distribution of potential cell therapies, but also a detailed view of their presence within different organs. Importantly, by using this imaging platform, we show that the route of administration affects the range of organs that the cells can reach, and, particularly, their propensity to form tumours. Our assessment suggests that cells are short-lived irrespective of whether they are administered via the venous or arterial circulation, and that the risk of cell persistence or tumour formation is dependent on the cell type, route of administration and immune status of the host. Crucially, we show that clinically used, human umbilical cord-derived mesenchymal stem/stromal cells form transient unexpected self-limited proliferations in various anatomical regions when administrated intravenously. The implications of this observation require further investigations and should be taken into account when clinical trials are considered.

## List of Abbreviations

ANOVA: analysis of variance
BLI: bioluminescence imaging
FACS: fluorescence activated cell sorting
GNRs: gold nanorods
hBM-MSCs: human bone marrow-derived mesenchymal stem/stromal cells
hKCs: human kidney cells
hUC-MSCs: human umbilical cord-derived mesenchymal stem/stromal cells
H&E: haematoxylin and eosin
IC: intracardiac
IV: intravenous
luc: luciferase (from firefly)
MRI: magnetic resonance imaging
MSCs: mesenchymal stem/stromal cells
mKSCs: mouse kidney-derived stem cells
mMSCs: mouse mesenchymal stem/stromal cells (D1 line)
MOI: multiplicity of infection
MSOT: multispectral optoacoustic tomography
PBS: phosphate buffered saline
PCV: packed cell volume
RMTs: regenerative medicine therapies
ROI: region of interest
SCID: severe combined immunodeficient
SPIONs: superparamagnetic iron oxide nanoparticles

## Declarations

### Ethics approval and consent to participate

All animal experiments were performed under a licence granted under the revised UK Animals (Scientific Procedures) Act 1986 and were approved by the University of Liverpool ethics committee.

## Consent for publication

Not applicable

## Availability of data and materials

The datasets supporting the conclusions of this article are included within the article and its additional files.

## Author Contributions

L.S., A.T., J.Sh., R.H., L.B., C.A.: conception and design, acquisition and/or assembly of data, data analysis and interpretation, manuscript writing, final approval of manuscript M.B., J.C., I.S., L.R.: acquisition and/or assembly of data, final approval of manuscript J.Sm., E.A.: provision of study material, final approval of manuscript C.H., R.L., M.J.R., D.J.A., H.P., B.K.P.: conception and design, data analysis and interpretation, final approval of manuscript P.M., B.W.: conception and design, data analysis and interpretation, manuscript writing, final approval of manuscript

## Acknowledgements

The authors would like to thank Carmel Moran and Adrian Thomson, University of Edinburgh, Tammy Kalber and Daniel Stuckey University College London, and Aleksandra Rak-Raszewska, for providing help and technical guidance in optimising the US-IC injection method, and Thomas Wilm, University of Liverpool, for help with the confocal imaging. All *in vivo* imaging was carried out in the Centre for Preclinical Imaging, University of Liverpool

